# A unique malignant cell type per patient tumor is encoded in each cancer cell transcriptome

**DOI:** 10.1101/2024.10.30.620989

**Authors:** Mirca S. Saurty-Seerunghen, Elias A. El-Habr, Yossi Eliaz, Léa Bellenger, Christophe Antoniewski, Hervé Chneiweiss, Marie-Pierre Junier

## Abstract

Deciphering shared features between patients through unsupervised analyses of tumor single-cell transcriptomes is hindered by the predominant clustering of malignant cells based on the patient’s tumor of origin. In contrast, cancer-associated non-malignant cells within tumors cluster according to their somatic cell type (e.g. macrophage, fibroblast, astrocyte), regardless of the individual patient. We investigated the origin of this contrasting clustering behavior using computational analyses and novel data sampling techniques across 160 tumors representing 9 distinct cancer types. This led to the unexpected disclosing of tumor-specific malignant cell types. We demonstrate that tumor-driven malignant cell clustering is independent from technical or computational biases, nor is it reducible to defined gene sets with tumor-specific expression across all examined cancers. On the opposite, we unveil redundant information dispersed across the transcriptome that encodes a unique identity shared by the malignant cells within each tumor. Additionally, we found a similar genomic encoding for the identity of normal cell types. Finally, we demonstrate that malignant cell identities are maintained across space and over time, as are the identities of normal somatic cells types. These findings suggest the establishment of a distinct type of malignant cells within each tumor, robustly and diffusively encoded across the entire transcribed genome.

## 1. Introduction

One of the main challenges of cancer research is the identification of common patient characteristics, necessary both to understand the developmental dynamics of a given tumor type and to allow the development of therapies that will benefit all patients. The development of single-cell RNA-sequencing (scRNA-seq) technologies has provided a unique opportunity to delve into the complexity of tumors in the real-life context of patients. This advancement has led to significant progress in understanding intra-tumor heterogeneity, including the diversity of normal cell types intermingled with malignant cells (Ho et al. 2018; Hong et al. 2019; Suva and Tirosh 2020; Ren et al. 2021). These findings have primarily relied on unsupervised clustering analyses conducted after filtering the datasets to keep only features considered to be most informative of the data structure, such as highly variable genes (HVGs) (Darmanis et al. 2017) or most expressed genes (Patel et al. 2014; Tirosh et al. 2016a; Tirosh et al. 2016b; Venteicher et al. 2017; Filbin et al. 2018). Strikingly, unsupervised clustering analyses of scRNA-seq datasets from both solid and non-solid tumors consistently result in a predominant grouping of malignant cells according to their tumor of origin, while normal cells from the same tumors are mainly grouped by somatic cell type (Tirosh et al. 2016a; Tirosh et al. 2016b; Chung et al. 2017; Darmanis et al. 2017; Puram et al. 2017; Filbin et al. 2018; Lambrechts et al. 2018; Jang et al. 2019; Ma et al. 2019; Saurty-Seerunghen et al. 2019; Chen et al. 2020; Maynard et al. 2020; Wiseman et al. 2020; Petti et al. 2022). These consistent results emerge irrespective of the features selected for analysis or the clustering algorithm used. Answers to this limitation have been provided by analyzing data tumor per tumor and then identifying commonalities between the results obtained (Tirosh et al. 2016b; Yuan et al. 2018; van Galen et al. 2019), or by merging data from different tumors and analyzing them as a whole after standardization (i.e. subtracting from each expression value the gene expression mean and dividing by its standard deviation across cells within a given tumor) (Muller et al. 2016), or by circumscribing cell groups according to a common signature known to be associated with a developmental-like cell state or a cell behavior (Neftel et al. 2019; Ren et al. 2021; Wu et al. 2021; Saurty-Seerunghen et al. 2022). These solutions have been implemented in the context of the assumption that tumor-driven malignant clustering is linked to technical variations in the sequencing between tumors (referred to as the batch effect) or to the tumor genomic specificities, notably chromosome copy number variations (CNV) that markedly impact gene expression levels (Davis-Marcisak et al. 2021; Vazquez-Garcia et al. 2022; Zhang et al. 2022).

Yet, the dependence of malignant cell clustering on the tumor of origin can hardly be attributed to scRNA-seq technical biases, since normal cells cluster independently from their tumor of origin. A simple dependence on tumor genomic specificities is not entirely obvious either, since distinct genomic clones coexist in tumors (McGranahan and Swanton 2017). Consistently, we previously noted that excluding CNV-associated gene values from the analysis of a scRNA-seq dataset of four patients’ glioblastoma did not alter the predominant clustering of malignant cells according to the tumor from which they originated (Saurty-Seerunghen et al. 2019). This ensemble of observations points to a gap in our understanding of the specificities of scRNA-seq analysis, and hence in its exploitation for the patients’ benefit.

This led us to undertake a formal exploration of the origin of this tumor-driven clustering of malignant cells. To that aim, we applied robust complementary computational analyses combined with original modes of data sampling to large datasets encompassing different types of solid tumors affecting the brain, the skin, the ovaries and the head and neck, and extended the analysis to normal cells. Our findings demonstrate that tumor-driven malignant cell grouping is unrelated to tumor genomic specificities in all cancers analyzed. Interrogating the structure of the information that distinguish the ensemble of malignant cells of each given tumor revealed its dispersion across the whole transcriptome of each malignant cell. Analyses of normal cell transcriptomes demonstrate a likewise structure of the information driving normal cell clustering according to their somatic cell type, and additionally shows that information linked to somatic cell types overshadows that which could distinguish one human individual from another. Finally, we demonstrate that the information characterizing malignant cells from a given tumor is maintained across space and over time, as is the identity of normal somatic cell types. Altogether, these results point to the establishment of a single type of malignant cells unique to each tumor, which set each tumor apart even among tumors belonging to the same class of cancer.

## Results

### Divergent impact of patient tumors on malignant and non-malignant cell grouping

Unsupervised grouping analysis using all detected genes consistently results in a tumor-driven grouping of malignant cells across all studied cancers, as illustrated with Uniform Manifold Approximation and Projection (UMAP) and alluvial plot representations for a scRNA-seq dataset (Darmanis et al. 2017) obtained from 4 patients’ tumors (Fig. 1A). On the opposite, the same analysis applied to non-malignant cells obtained from the same four tumors results in clusters populated by cells coming from distinct tumors and dominated by somatic cell types (Fig. 1B). We extended unsupervised grouping analysis to a total of fourteen datasets from brain and non-brain cancers obtained with differing sequencing techniques (Supplemental Table S1). We used Normalized Mutual

**Figure 1.**
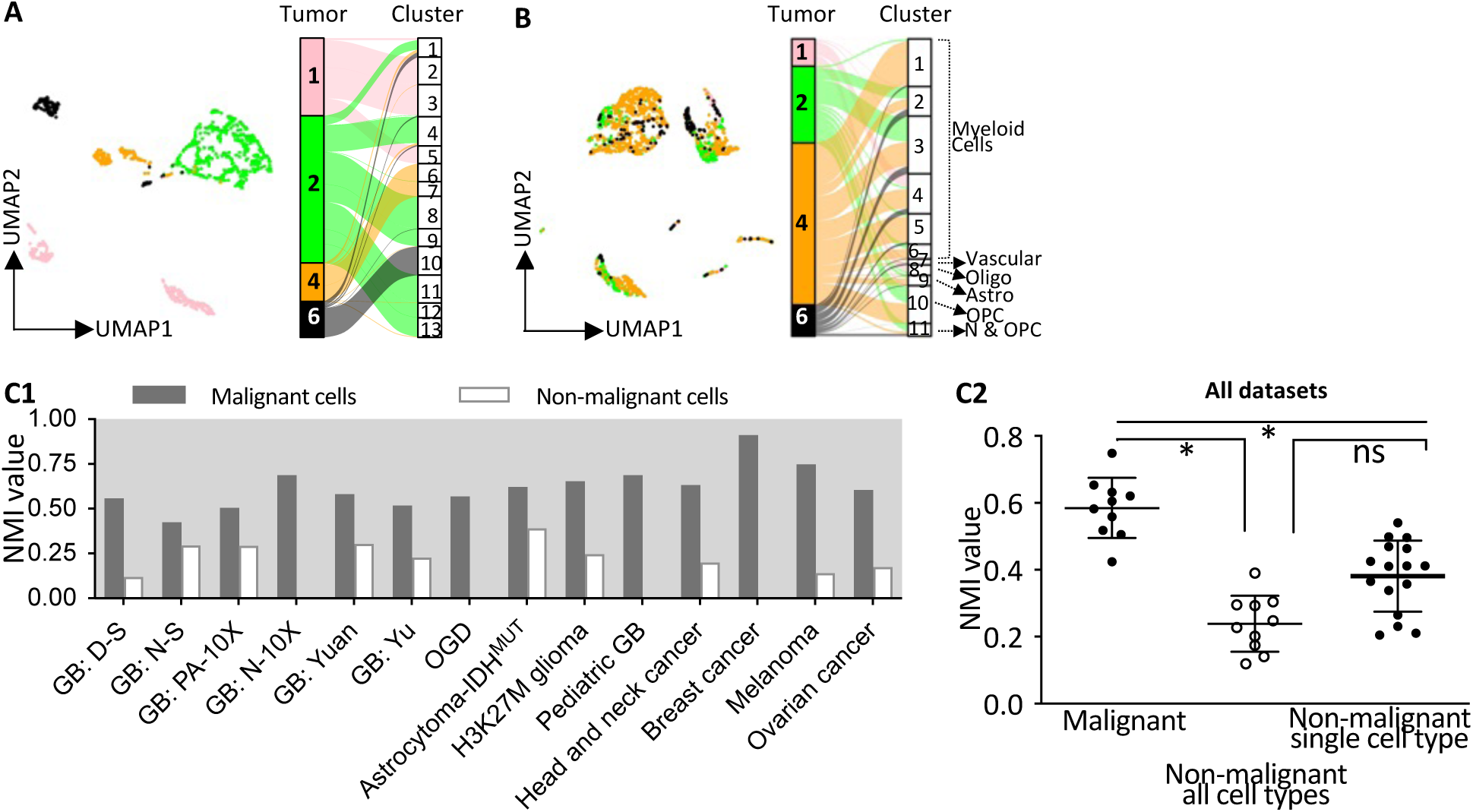
Tumor-driven malignant cell grouping in all types of cancers. **A. Tumor-driven malignant cell grouping** illustrated with the example of the Darmanis and colleagues dataset containing scRNA-seq from 4 glioblastoma. Left panel: UMAP representation with malignant cells colored according to the tumor from which they were harvested. Right panel: Alluvial plot representation showing the repartition of cells from each tumor (colored by tumor) between the clusters resulting from unsupervised grouping analysis. **B. Independence of non-malignant cell grouping from the tumor from which they are harvested** illustrated with the example of the Darmanis and colleagues dataset from 4 patients’ tumors. The somatic cell types represented in each cluster are annotated on the alluvial plot. Oligo, oligodendrocytes; Astro, astrocytes; OPC, Oligodendrocyte progenitor cells; N, neurons. **C. Tumor contribution to cell clusters.** Each Normalized Mutual Information (NMI) value corresponds to the grouping based on expression of all detected genes of malignant, and when available non-malignant cells. **c1:** Bar plot representation of NMI values for individual datasets. GB: glioblastoma with wild-type form of the Isocitrate dehydrogenase (IDH) genes. IDH^MUT^: mutant form of IDH1 gene. OGD: IDH1^MUT^ 1p/19q-codeleted Oligodendroglioma. See text for further details. **c2:** Dot plot illustrates reduced NMI values of non-malignant cell grouping compared to NMI values of malignant cells across all datasets analyzed. NMI values of non-malignant cells corresponding to unsupervised clustering of cells of different somatic types (Non-malignant all cell types) and of cells of a single type (Non-malignant single cell type). Kruskal-Wallis test, Mean ± SD, *: p <0.0001 (Malignant cells vs Non-malignant all cell types), p = 0.0057 (Malignant cells vs Non-malignant single cell type).

Information (NMI) to assess each tumor contribution to each cluster of malignant and non-malignant cells. Highest NMI values were systematically obtained for malignant cell clustering, indicating that cells predominantly group according to the tumor from which they are harvested (Fig. 1C). In contrast, non-malignant cell clustering was characterized by lower NMI values (Fig. 1C). We envisaged the possibility that technical variations might differentially affect non-malignant and malignant cells, or that information related to normal cell types could mask that related to the patients. NMI values resulting from clustering analyses performed on ensemble of normal cells of a given somatic cell type, remained lower than the ones obtained with malignant cell clustering (Fig. 1C2). These results confirm that the grouping of non-malignant cell is independent from the patient tumor. They also suggest that information related to somatic cell type is a stronger driver of unsupervised cell clustering than information specific to each human individual. This distinct clustering behavior between malignant and non-malignant cells from the same tissue indicates that factors external to the cells, such as technical biases, cannot account for the tumor-driven grouping of malignant cells.

### Independence of tumor-driven malignant cell grouping from computational biases and tumor-specific gene expression features

Although the contrasting grouping observed between malignant and non-malignant cells rules out technical biases as responsible for tumor-driven malignant cell grouping, we considered them for the sake of consistency of our exploration. Technical variations like differences in sample processing, differences in sequencing depth, cell lysis and reverse transcription efficiency were minimized through customary normalization of raw counts by sequencing library size. Another major source of technical bias is the variations in sample-dependent gene detection failures (referred to as dropouts). In scRNA-seq experiments, low amounts of RNA molecules in individual cells, inefficient mRNA capture, stochastic mRNA expression and amplification failures result in transcripts being missed, and consequently in cell-to-cell variability in transcriptome coverage (Qiu 2020). We tested the impact of dropouts by inferring and imputing the “missing” expression values. As expected, NMI values of malignant cell grouping after dropout imputation were not reduced compared to NMI values of grouping performed without data manipulation (Fig. 2A-D, orange-framed bars, Supplemental Table S2). They were rather increased in a statistically significant manner when considering the ensemble of all cancer datasets analyzed (Fig. 2E). This increase is consistent with the reliance of dropout imputation on information of the same gene expression borrowed from cells with global transcript profiles most close to the one of the cells exhibiting the dropout (Li and Li 2018).

**Figure 2.**
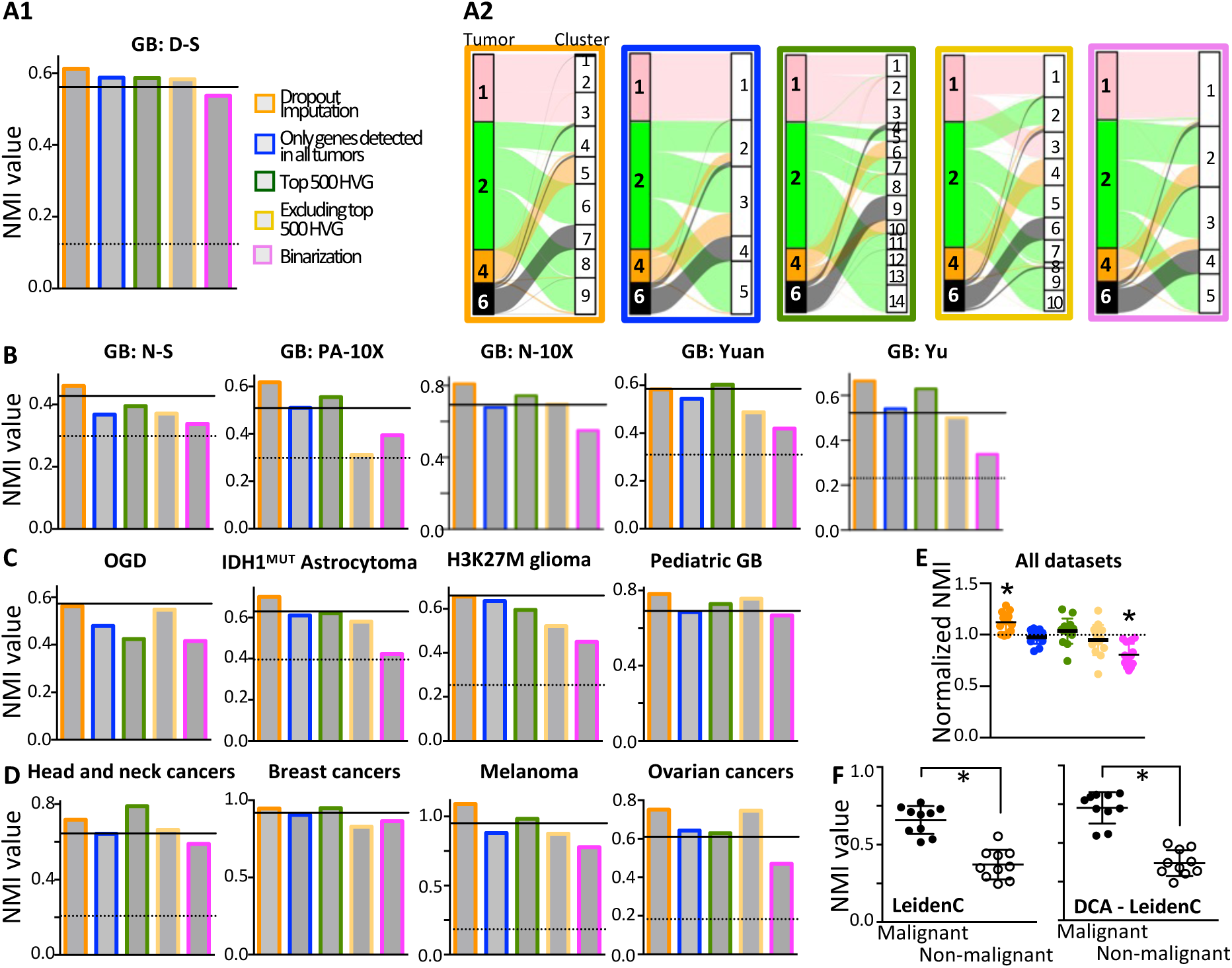
Tumor-driven malignant cell grouping is independent of technical bias or tumor-specific characteristics in gene expression for all cancers studied. Bar plots illustrate Normalized Mutual Information (NMI) values obtained after data manipulation i.e. following either dropout imputation (orange outline), using only genes detected in all tumors (blue outline), using the 500 most highly variable genes (HVG, green outline), as well as following exclusion of the 500 most variable genes (yellow outline), or following binarization of gene expression (pink outline). Horizontal solid and dotted lines correspond to the NMI values obtained from analyses performed without data manipulation of malignant and when available non-malignant cells, respectively. **A.** GB malignant cells from the scRNA-seq dataset of Darmanis and colleagues (2017). Bar plot (a1) and alluvial plot (a2) representations of the contribution of each tumor to each cluster following data manipulations. Compare with alluvial plot of Fig. 1A. **B.** GB malignant cells from independent datasets obtained with either SMART-seq2 (N-S), 10X Genomics single-cell RNA-seq (N-10X, PA-10X), or in-house sequencing technologies (Yuan, Yu). **C.** Malignant cells from other classes of brain tumors of the adult (OGD and IDH1^MUT^ Astrocytoma) and the infant (H3K27M glioma diffuse gliomas and pediatric gliomas). IDH1^MUT^: isocitrate dehydrogenase 1, mutated form. OGD: IDH1^MUT^ 1p/19q-codeleted Oligodendroglioma. SMART-seq2 technology. **D.** Malignant cells from non-brain cancers including head and neck cancers (SMART-seq2 sequencing technology), primary breast cancers (Fluidigm C1 technology), melanoma (SMART-seq2 technology) and ovarian cancers (10X Genomics technology). **E.** Tumor-driven cancer cell grouping consistently decreases only following binarization of gene expression. NMI values presented with respect to the NMI value obtained without data manipulation for malignant cell grouping of each dataset (Normalized NMI). Dotted line marks the NMI value without data manipulation. One sample t-test, Mean ± SD, p<0.0003 (dropout imputation) and p<0.0001 (binarization). **F.** Reduced NMI values of non-malignant cell grouping compared to NMI values of malignant cells observed using two other analytical approaches. Dot plot illustrates reduced NMI values of non-malignant cell grouping compared to NMI values of malignant cells across all datasets analyzed. Compare with Fig. 1C2. Mann-Whitney, Mean ± SD, *: p<0.0001. M: Malignant cells, N: Non-malignant cells. See Fig. S2B for corresponding bar plot.

To evaluate the influence of tumor-specific biological features on malignant cell clustering, we first considered only the genes with the highest expression variability across all cells of each dataset (highly variable genes or HVG) because they reflect inter-tumor genomic variations with the largest impact on the global transcriptome (e.g. mutations, CNV). Including or excluding the 500 most HVG did not consistently modify the NMI values (Fig. 2A-D, green- and yellow-framed bars, Fig. 2E, Supplemental Table S2). All information on variability in gene expression levels was then eliminated by considering either only the genes detected in all tumors or by performing data binarization, where detected genes are assigned a value of one and non-detected genes a zero value. Cell clustering based on genes universally detected in all tumors of a given dataset yielded NMI values similar to those obtained with the entire transcript repertoire (Fig. 2A-D, blue-framed bars, Fig. 2E, Supplemental Table S2). Data binarization reduced but did not eliminate the influence of the tumor on malignant cell grouping (Fig. 2A-D, pink-framed bars, Fig. 2E, Supplemental Table S2). These results indicate that tumor-specific gene repertoires, whether in terms of detection or expression levels, are not sufficient to account per se for tumor-driven cell clustering. Finally, we investigated whether tumor-dependent cell grouping might be linked to the analytical pipeline (Supplemental Fig. S1A). Use of a different clustering method based on community detection (Leiden) yielded similar results, malignant cells having significantly higher NMI values than non-malignant cells (Fig. 2F, Supplemental Fig. S1B, Supplemental Table S2). Specific tumor-driven clustering of malignant cells was also observed when using deep count autoencoder (DCA) as an alternative data reduction approach for principal component analysis (PCA) (Fig. 2F, Supplemental Fig. S1B, Supplemental Table S2).

These findings collectively demonstrate that neither technical biases nor tumor-specific gene repertoires, whether in terms of detection or expression levels, are sufficient to explain tumor-driven cell clustering. They suggest that the clustering of malignant cells by tumors arises from information embedded within the transcriptome of all malignant cell constituting a tumor, which encode a unique identity for each tumor.

### Information encoding each tumor identity is scattered across cell transcriptomes

The persistent influence of the tumor on malignant cell grouping, despite extensive data manipulations and different analytical approaches, suggested that this influence cannot be reduced to inter-tumor specificities in gene expression or to variations in technical processing. This led us to consider that, for all cancer classes examined, information pertaining to a tumor influence on malignant cell clustering is dispersed throughout the cell transcriptomes. If this is indeed the case, we expect to detect it as long as random gene sampling covers a significant fraction of the transcriptome, then lose it with further reductions in the number of genes sampled, as schematically illustrated in Fig. 3A. Accordingly, NMI values of the clustering should vary based on the sampling of the dispersed information expected to encode the identity of the tumors. To test this hypothesis, we performed unsupervised clustering analyses based on progressively smaller numbers of randomly selected genes. For all cancer classes, we observed progressive decreases in NMI values of the clustering of malignant cells as the number of randomly-selected genes considered in the analyses decreased (Fig. 3B, Supplemental Fig. S2). The influence of the tumor was suppressed only when reducing the number of analyzed genes below 100 to 50, as indicated by the stabilization of NMI values. In other words, combined expression of over 100 genes is sufficient to retrieve the tumor identity for all cancers studied. Of note, the gene lists considered in these analyses were dissimilar from one another, with less than 30% overlap for gene set sizes of 250-5000 and almost no overlap for gene set sizes inferior or equal to 100 (Supplemental Fig. S3).

**Figure 3.**
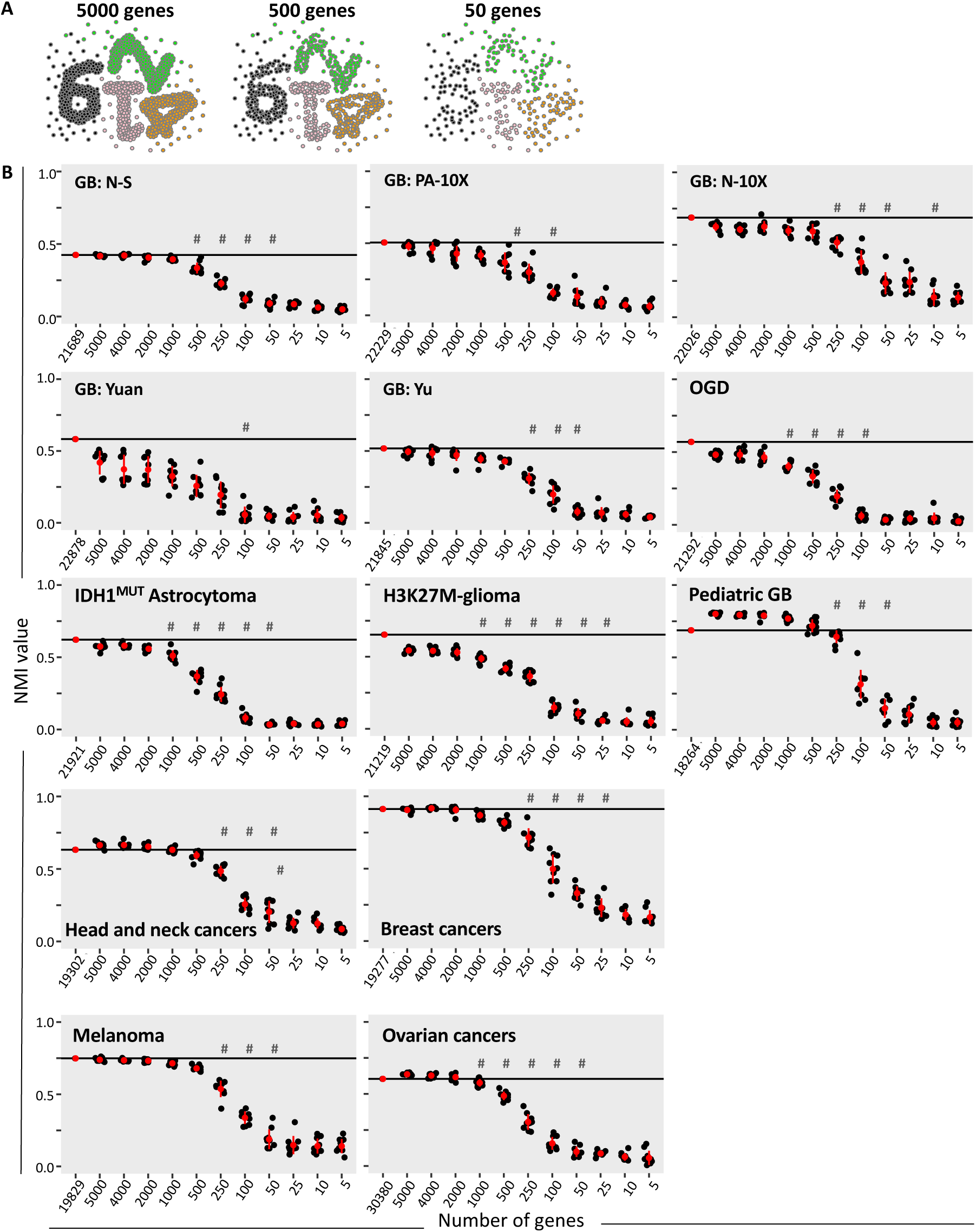
Tumor-specific identities of malignant cells are encoded by information dispersed through each cell transcriptome. **A.** Illustration of the hypothesis where each dot symbolizes a gene, and the number 1246 symbolizes the tumor-specific identity of malignant cells. An identity encoded by information dispersed through the whole cell transcript repertoire, is expected to be lost when including decreased number of randomly selected genes in unsupervised clustering analyses. **B.** Combined expression of over 100 genes is sufficient to retrieve a tumor-specific identity. Decreased NMI values with inclusion of decreased number of randomly selected genes in the grouping analyses in all scRNA-seq analyzed, regardless of the class of cancer considered or the sequencing technology implemented. Ten independent analyses performed with randomly selected genes for each gene number analyzed. Mean ± SD shown in red. #: statistically significant differences with the immediately prior sampling, Holm-Sidak’s multiple comparison test, adj-p value<0.05.

These results demonstrate that, irrespective of the class of cancer examined, all malignant cells from a given tumor share a common identity, which is encoded by information dispersed throughout the cells’ transcriptome.

### Malignant cell types defined by tumor identities that persist across space and over time

The genome-wide dispersion of the information specific to each tumor led us to envisage that it distinguishes malignant cells according to tumor-specific malignant cell types, as normal cells can be distinguished in unsupervised analyses on the basis of the ontogenically defined specific somatic cell type to which they belong. We tested this possibility by first determining whether normal cell types are also encoded by information dispersed throughout their transcriptomes. Unsupervised analyses were applied on scRNA-seq data of normal cells from non-cancerous brain samples and healthy bone marrow donors, as well as from non-malignant cells harvested from tumors. The results revealed a progressive loss of the predominance of somatic cell type on the clustering of normal cells as the number of randomly-selected genes included in the analyses was reduced, as evidenced by the reduced NMI values (Fig. 4, Supplemental Fig. S4). These results demonstrated that the structure of the information encoding normal somatic cell types is similar to that shared by malignant cells of a given tumor. We then determined whether information related to tumor-specific identity of malignant cells is, like information related to normal somatic cell types, maintained across space and over time. In other words, can we distinguish the tumor identity of a malignant cell independently of its spatial localization and the time at which it is scrutinized as we can distinguish, for example, a lymphocyte from other cell types regardless of the age of the individual and the organ where it is found? The tumor micro-environment can vary across different anatomic locations in terms of oxygen pressure, blood vessel density, growth factors, nutrients, metabolites or cellular composition among others (Hambardzumyan and Bergers 2015; Schiffer et al. 2018; Bikfalvi et al. 2023). It also evolves over time, according to natural tumor growth as well as in response to therapeutic assaults (Junttila and de Sauvage 2013). These varying environments are known to differentially impact the dynamics of gene expression in malignant cells. To determine whether malignant cells sampled at different locations of a patient’s tumor maintain a common identity, we analyzed two scRNA-seq datasets containing malignant cells from multiple territories of the same glioblastoma (Darmanis et al. 2017; Yu et al. 2020). This brain cancer is well known for its heterogeneous micro-environments (Bikfalvi et al. 2023). We also examined the robustness of the tumor identity across time using three datasets containing malignant cells from basal cell carcinoma, glioblastoma, or ovarian carcinoma sampled before and after patients’ treatment (Yost et al. 2019; Wang et al. 2022; Zhang et al. 2022). Unsupervised analyses were performed using all genes detected in each ensemble of cells from all territories or time points. Clustering results showed that glioblastoma cells from a given patient grouped together irrespective of their location within the tumor, as highlighted on UMAP (Fig. 5A, Supplemental Fig. S5A and B). Likewise, we observed a predominant grouping of ovarian carcinoma cells (Fig. 5B, Supplemental Fig. S5D), basal carcinoma cells (Supplemental Fig. S5C), or glioblastoma cells (Supplemental Fig. S6) based on their tumor of origin, regardless of their sampling prior or after patient treatment.

**Figure 4.**
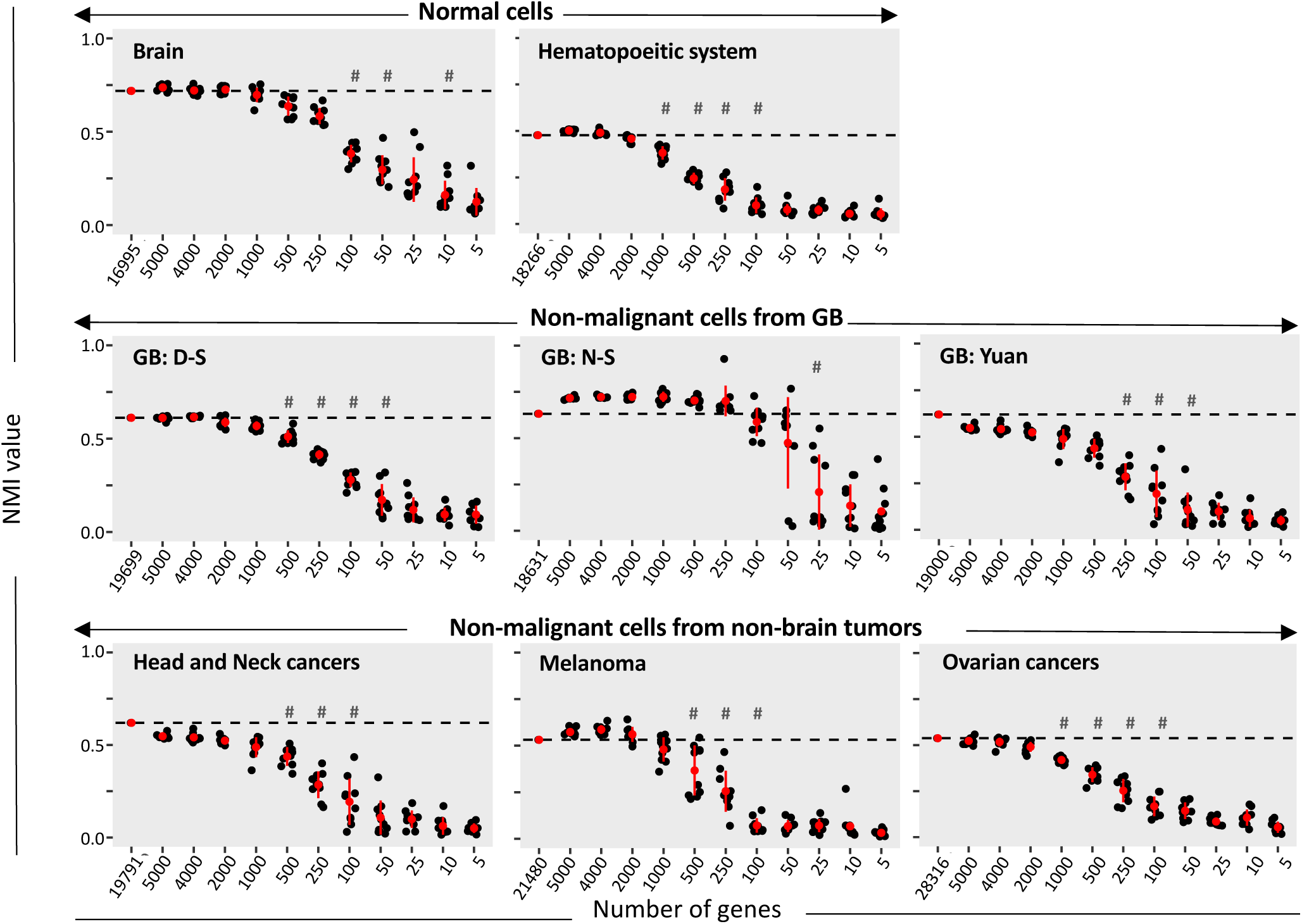
Non-malignant cell types are also encoded by information dispersed through the transcriptome of each cell. Down-sampling gene numbers decreases the influence of the somatic cell type on the grouping of normal cells. Datasets analyzed include hematopoietic cells (myeloid, erythroid and lymphoid), epithelial cells, neural cells (oligodendrocytes, astrocytes, neurons), vascular cells, fibroblasts, and myocytes. NMI values are depicted. Ten independent analyses performed with randomly-selected genes for each gene number analyzed. Mean ± SD shown in red. #: statistically significant differences with the immediately prior sampling, Holm-Sidak’s multiple comparison test, adj-p value<0.05.

**Figure 5.**
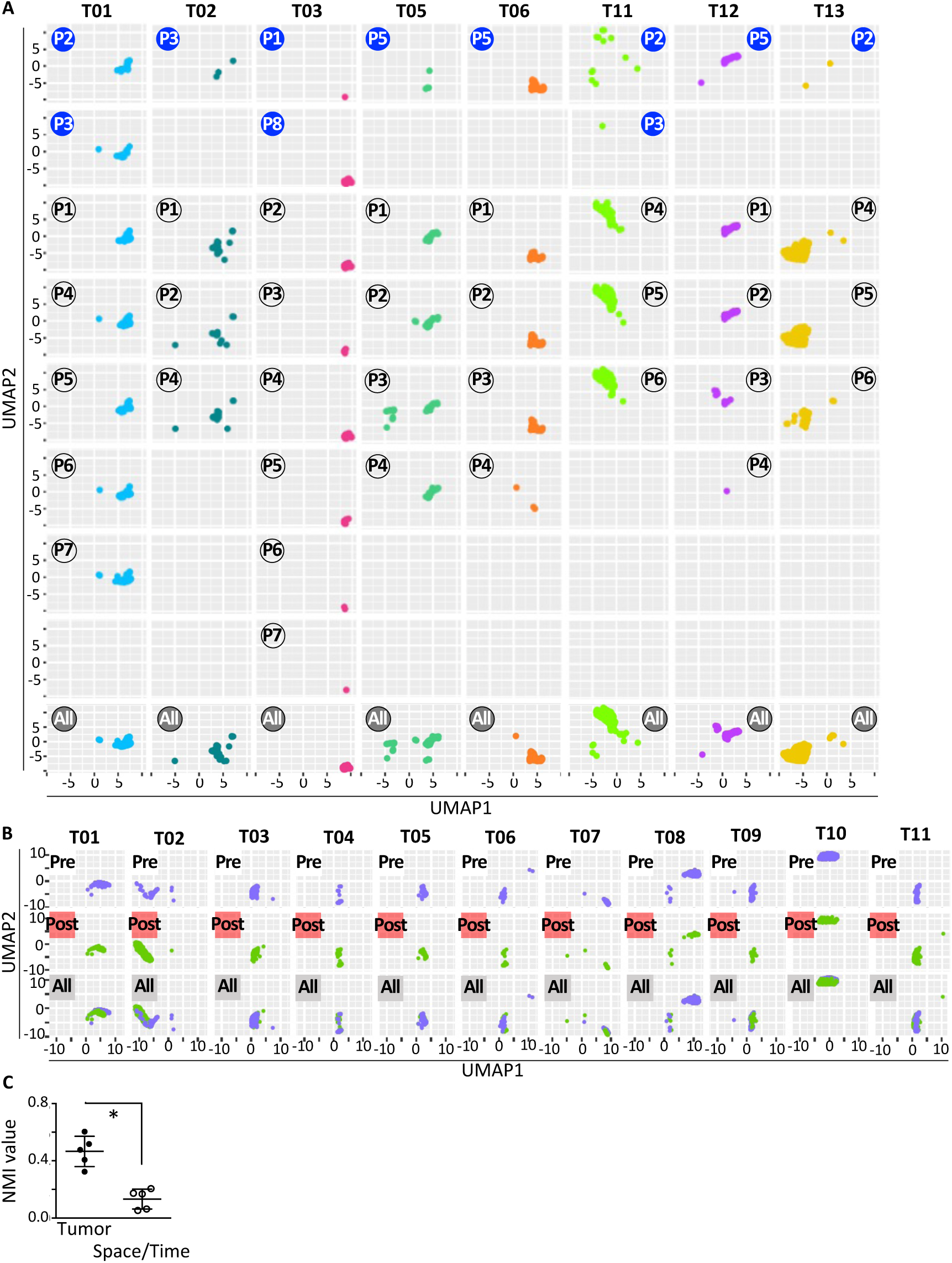
Tumor-specific identities of malignant cells are maintained across varying spatial environments and over time. **A-B.** UMAP representations with cells colored according to the tumor (T) from which they were collected. **A.** Malignant cells collected from different glioblastoma (GB) regions group together by tumor of origin, irrespective of the tumor region they come from. GB scRNA-seq datasets from Yu et al 2020. The numbers of sampled regions are circled. Blue circles correspond to malignant cells sampled from the peri-tumoral brain, black circles to cells from the tumor, and grey circles to cells from both regions. **B.** Malignant cells collected from ovarian cancer samples group together by tumor of origin, irrespective of the treatment time point at which they were collected. scRNA-seq dataset from Zhang et al et al 2022. White squares correspond to cells collected prior treatment, red squares to cells collected post-treatment and grey squares to cells from both time points. **C.** Reduced Normalized Mutual Information (NMI) values corresponding to the contribution of microenvironments and treatment time points to cell clusters (Space/Time) compared to the tumor contribution to cell clusters (Tumor). Mann-Whitney test, Mean ± SD, *: p <0.0079, n=5 independent scRNA-seq datasets. See text for further details.

In a coherent manner, all NMI values calculated for the contribution of microenvironments or treatment time points to cell clusters were lower than the ones calculated for the tumor contribution to cell clusters (Fig. 5C, Supplemental Table S2).

These results demonstrate the conservation of information related to the tumor identity, regardless of changes in tumor micro-environments or of tumor evolution over time. Together with our finding that tumor identity and normal somatic cell types are encoded likewise by genome-wide dispersed information, they support the view that the tumor identities highlighted by our analyses specify types of malignant lineages, unique to each patient tumor.

## Discussion

Our study, exploring the source of the contrasting grouping of normal and malignant single cell transcriptomes, resulted in demonstrating that a given patient tumor is composed of malignant cells belonging to a single broad cell type specific to each tumor. Altogether, our findings demonstrate that irrespective of the type of cancer considered, all malignant cells of a given tumor share common information dispersed throughout the cells’ transcriptome that constitute a unique identity, stable across space and over time. This identity is unique to each tumor since it suffices to distinguish its malignant cell populations from those of other tumors of its kin, even when considering cancers whose classification is refined by knowledge of the founding or most prevalent mutation(s). In other words, if tumors of the same cancer class contained similar malignant cell types, they would group together in cross-tumor comparisons, as observed for non-malignant cells populating the tumors or for normal cells across healthy individuals. This is a generalizable property of cancer cells as illustrated in our analyses of the dataset of H3K27 mutated pediatric glioma (Filbin et al. 2018) or of the datasets of IDH mutated diffuse glioma (Tirosh et al. 2016b; Venteicher et al. 2017), and is visible as well on previously published UMAPs showing tumor-driven clustering of malignant cells among ER+, HER2+ or TNBC breast cancer sub-types (see figure 1d-f in (Wu et al. 2021)).

The two most obvious candidates that could explain tumor-driven cell grouping - technical biases and tumor molecular features-do not account for transcriptome characteristics underlying this mode of grouping. Technical biases are excluded since clustering of the non-malignant cells populating the tumors escape the influence of the tumors, as shown here as well as in all previously published analyses of scRNA-seq derived from tumors (e.g. (Tirosh et al. 2016a; Puram et al. 2017; Lambrechts et al. 2018; Ma et al. 2019; Saurty-Seerunghen et al. 2019; Chen et al. 2020; Maynard et al. 2020; Petti et al. 2022)). Of note, non-malignant cells populating the tumors group independently of their tumor of origin, despite their deviation from their healthy counterparts linked to the pathological context. Accordingly, our comparative analyses of non-malignant cells from different somatic cell types or from a unique somatic type show that the dominating factor driving their clustering corresponds to the somatic cell type rather than the patient identity. We also verified that another possible technical bias - computational methods used-is not involved, since differing methods yield similar tumor-driven grouping of malignant cells. Salient molecular features of tumors constituted by genomic alterations are the other obvious candidate. Our analyses performed after excluding tumor-specific features provide a formal demonstration that tumor-driven cell grouping cannot be reduced to tumor-specific gene repertoires. Of note, all tumors we analyzed contain multiple genomic cell clones as shown by CNV analyses. The demonstration that tumor-specific genomic features are insufficient to account for tumor-driven malignant cell grouping led us to uncover an unexpected structuring of the information related to malignant cell identity. We found that this identity is encoded by information dispersed throughout the transcriptome of each malignant cell within a given tumor, and that it is highly redundant, with 100-1000 genes being sufficient to reveal this identity. Remarkably, we found a similar encoding of information pertaining to the identity of normal somatic cell types. These results provided the first evidence that the malignant cell identities disclosed by our analyses define tumor-specific malignant cell types.

Establishment of atlases of cell types in different organisms, organs and developmental stages has provided strong support for the biological relevance of unsupervised clustering of single cell transcriptomes to categorize normal cells into cell types (Zeng 2022). Importantly, these works show that somatic cell types can be uncovered from single cell transcriptomes in a consistent manner regardless of the cell context, indicating that somatic cell type identity encoded in transcriptomes dominates over variations in molecular correlates linked to different geographical or temporal contexts encountered by the cell, as well as to the genomic background of the individuals. Coherently, we observed a remarkable stability of the information pertaining to malignant cell identities across diverse spatial and temporal contexts. Glioblastoma are highly infiltrative tumors that disperse cells through the brain parenchyma. They thus encompass two totally distinct micro-environments in terms of cellular composition: the tumor periphery dominated by normal cells, and the tumor core dominated by malignant cells. Leveraging datasets with malignant cells sampled from these distinct regions, we demonstrate that tumor-driven cell clustering remains consistent, irrespective of the environment from which the cells are harvested. The maintenance of malignant cell identity despite environmental changes is also evidenced by the common clustering of malignant cells from primary tumors and their metastases in different organs (Puram et al. 2017; Quah et al. 2023). To assess the stability of this identity over time, we utilized longitudinal tumor samples encompassing primary tumor specimens and those collected from tumor recurrences after therapy. In the three cancer types examined (ovarian cancer, basal cell carcinoma and glioblastoma), our results show that malignant cells from pre- and post-treatment stages cluster together based on the tumor of origin. Taken together, our analyses of malignant cells from distinct tumor regions or timepoints demonstrate an exceptional robustness of malignant cell identities, all the more remarkable considering the well-known evolution of malignant cells in response to therapeutic pressures (Marine et al. 2020; Tyner et al. 2022). Importantly, they support the establishment of a unique malignant cell type per tumor despite the diverse environmental pressures and genomic alterations encountered within each tumor.

The tumor-specific type of malignant cells highlighted in this study are likely to be divisible into sub-types, considering that the strength of its impregnation eclipses the imprints of other biological information. Failure to group malignant cells from different tumors despite their sharing of similar major genomic alterations or their derivation from the same type of founding cells is a notable illustration of this strength (Sottoriva et al. 2013; Wang et al. 2014; Lee et al. 2017; Neftel et al. 2019; Couturier et al. 2020; Wu et al. 2021). It also prevents malignant cells from grouping based on their most biologically relevant feature, their functional states, i.e. what they do at the time of their harvest. Malignant cells displaying variable expression levels of genes defining normal somatic cell types have been highlighted in each tumor using cell type-specific molecular signatures (Couturier et al. 2020; Wu et al. 2021) or supervised clustering based on molecular signatures extracted from the datasets under scrutiny (Neftel et al. 2019). The interpretation of these findings in relation with tumor growth, hierarchical organization and developmental history however remains to be stabilized across tumors from patients with a similar type of cancer, and needs to be explored further. Likewise, cells assuming similar functions, such as proliferation or invasion, known to be present in each patient’s tumor, can only be potentially identified by calculating their expression scores of gene modules based on previously acquired experimental evidences or by supervised clustering analyses (Tirosh et al. 2016a; Saurty-Seerunghen et al. 2022). Importantly, these findings show that various phenotypic features can be extracted from the transcriptomes of cancerous cells. In face of these findings, detection of a single type of malignant cell per tumor can be envisaged as a reflect of the plasticity of cancerous cells that have to adapt to the constantly changing environment of a growing tumor, a process already recognized in some cancers (Kong et al. 2020; Huang et al. 2021; Yabo et al. 2021; Pérez-González et al. 2023). Such adaptations requiring the rapid tuning of the transcriptome through continuous variations in gene expressions are indeed expected to blur the boundaries between distinct cell sub-types.

The genome-wide distribution and the high redundancy of information related to somatic cell type highlighted by our results are coherent with the known complexity of the establishment of normal cell lineages which results from complex interplays between cell intrinsic and extrinsic signals. It is also coherent with the view that normal cell types are specified by core sets of transcription factors whose targets can be distributed throughout the genome (Arendt et al. 2016). Specification of the unique cell type identified in each tumor is also likely to result from a similar mixture of events, further complexified by the advent of stochastic genomic alterations, with its establishment and maintenance depending on self-propagating transcriptional states such as those driven by inheritable super-enhancers, clusters of DNA-regulatory elements controlling gene expression programs (Hnisz et al. 2013). Although the high redundancy of the information related to malignant cell types makes it challenging to identify the core molecular characteristics constituting a malignant cell type, their identification could lead to the pining down of actionable weaknesses shared by most of the malignant cells populating a given tumor. Multi-omics approaches integrating the epigenetic profile of the cells, such as derived from single cell ATAC-seq, DNA methylation or histone based-chromatin immunoprecipitation, may help achieving this goal, although each alone can also result in grouping cells per tumor as can be observed in unsupervised grouping of scATAC-seq profiles (Regner et al. 2021; Kumegawa et al. 2022).

In summary, our findings demonstrate for the first time that each patient’s tumor establishes a distinct and unique type of malignant cells, which is preserved across different environments, and enduring over time and through therapeutic pressure. The identity of the malignant cell type encoded in single-cell transcriptomes supersedes all other information, whether related to the functional state of the cell, its degree of differentiation or its genomic alterations. Single-cell transcriptomes thus offer a novel view of tumor identity based on the single malignant cell type established in each tumor, rather than it being primarily defined by genomic alterations.

## Methods

### Data acquisition

The single-cell transcriptomes were downloaded from public databases (Supplemental Table S1). Analyses were performed using independent single-cell RNA-seq datasets from adult glioblastoma (Darmanis et al. 2017; Yuan et al. 2018; Neftel et al. 2019; Yu et al. 2020; Pombo Antunes et al. 2021; Wang et al. 2022), from other adult-type diffuse gliomas including IDH1-mutant 1p/19q-codeleted oligodendroglioma (Tirosh et al. 2016b) and IDH1-mutant astrocytoma (Venteicher et al. 2017), from pediatric gliomas including pediatric glioblastoma (Neftel et al. 2019) and H3K27M-glioma (Filbin et al. 2018), and from non-brain tumors including melanoma (Tirosh et al. 2016a), basal cell carcinoma (Yost et al. 2019), head and neck cancers (Puram et al. 2017), breast cancers (Chung et al. 2017) and ovarian cancers (Zhang et al. 2022). We also used scRNA-seq datasets from non-cancerous brain samples (Darmanis et al. 2015) and healthy bone marrow donors (van Galen et al. 2019). All these datasets included cells coming from multiple human individuals.

We used log_2_-transformed Counts Per Million or Transcripts Per Million (log_2_(CPM+1) and log_2_(TPM+1) respectively) to allow comparison of read abundance across libraries of different sizes. CPM or TPM corresponds to the number of gene-mapped reads (Counts or Transcripts) divided by the total number of mapped reads per cell, the resulting number being then divided by one million (Per Million). This sequencing depth normalization is required for between-sample comparisons (i.e. when comparing gene expression between cells). TPM adds another level of transformation prior to sequencing depth normalization. In that case, the number of gene-mapped reads is divided by the transcript’s length so as to take into account the length of transcripts in the estimation of read counts. To avoid potential analytical bias due to scarcely detected genes, we filtered out genes detected in less than 3 cells prior to grouping analysis.

### Identification of malignant and non-malignant cells

Malignant and normal cells were distinguished as we previously described (Saurty-Seerunghen et al. 2022) either according to the cell annotations provided by the authors of the corresponding datasets, or when absent based on inference of copy-number variations (CNV), a hallmark of malignant cells. In brief, expression data in log2(CPM/100 + 1) values were processed using a three-step approach: CNV inference using CONICSmat R package (Muller et al. 2018), marker gene expression and unsupervised cell clustering. The default filtering and normalization procedures were followed for CNV inference, as outlined in https://github.com/diazlab/CONICS/wiki/Tutorial---CONICSmat;---Dataset:-SmartSeq2-scRNA-seq-of-Oligodendroglioma. For each region, CONICSmat likelihood ratio test adjusted p-value <0.001 and a difference in Bayesian Criterion >300 were retained as CNVs. The GMM-based CNV predictions were then used to group cells into potential malignant and non-malignant groups and visualized using heatmaps (ComplexHeatmap R package (Gu et al. 2016)) and Uniform Manifold Approximation and Projection (UMAP, umap R package) plots based on 500 most variable genes (Supplementary Fig. S7A-B). Second, expression of marker genes for pan-immune cells (PTPRC/CD45), macrophages (ITGAM, FCGR3A/CD16A, CD14), microglia (CSF1R, TMEM119), T-cells (CD2, CD3D) and oligodendrocytes (MOG, MAG) was highlighted on UMAP plots (Supplementary Fig. S7C). Finally, a hierarchical clustering followed by a K-means clustering on the UMAP components (FactoMineR R package (Husson et al. 2010)) was applied to identify cell groups that were most similar to one another (Supplementary Fig. S7D). Using CNV status predictions, marker gene expressions and the clustering result, the cells were marked as malignant when they harbored CNVs, clustered together on UMAP and were devoid of normal cell markers (Supplementary Fig. S7E).

### Unsupervised grouping analyses

All R and Python scripts used are provided in Supplementary files 1 and 2, respectively. Three approaches were used for cell grouping analyses using all detected genes (Supplementary Fig. S1A): Hierarchical Clustering on Principal Components (HCPC), principal component analysis (PCA) followed by Leiden Clustering without or with prior data denoising with deep count autoencoder (DCA) (hereafter designated as LeidenC and DCA-LeidenC, respectively).

The HCPC approach (Husson et al. 2010) combines the hierarchical and partitioning clustering strategies. The hierarchical clustering has the advantage of being one of the most robust clustering strategies – it yields consistent results with different number of clusters and shows little variation across datasets (Lu et al. 2019). The cut of the hierarchical tree is introduced as the initial partition of the K-means algorithm and several iterations of this algorithm are done to improve the cell grouping (Husson et al. 2010). HCPC was performed with FactoMineR package (version 2.4) as previously described (Saurty-Seerunghen et al. 2019). The number of clusters was determined by analyzing the dendrogram obtained upon hierarchical clustering of the data and the optimal number of clusters suggested by the Silhouette method (NbClust R package, version 3.0 (Charrad et al. 2014)). When the optimal number suggested was coherent with the aspect of the dendrogram, then the optimal number was used. Else, the number of clusters was decided using the dendrogram. UMAP was used to visualize transcriptional similarity between cells in a low dimension embedding (umap R package, version 0.2.10). Alluvial plots were generated to visually determine the contribution of each tumor to each cluster (ggplot2 R package, version 3.4.3). HCPC was used for all analyses, unless otherwise specified.

LeidenC was performed using Python. The normalized gene expression matrix was converted to a sparse matrix using the coo_matrix format from the SciPy library (version 1.15.1 (Virtanen et al. 2020)) to ensure efficient storage and manipulation. Dimensionality reduction was then performed using PCA from Scanpy python package (version 1.9.6 (Wolf et al. 2018)). The PCA embedding was then used for cell clustering using the Leiden algorithm (Scanpy package).

For DCA-LeidenC performed using Python, the normalized gene expression matrix was converted to a sparse matrix as for LeidenC. The data were then transformed using DCA (DCA python package, version 0.3.4 (Eraslan et al. 2019)) to aid in denoising the expression data and capturing the underlying structure. Finally, Leiden clustering was applied to the DCA-transformed data as described above.

### Data manipulation prior to cell clustering

Imputation of dropouts was performed using the scImpute R package (version 0.0.9) (Li and Li 2018) with the following parameters: input expression data in log_2_(CPM+1) or log_2_(TPM+1), drop_thre = 0.75, Kcluster = 1 and all other parameters set to default. Gene expression binarization and identification of genes detected in all tumors were implemented as previously described (Saurty-Seerunghen et al. 2019). 500 genes with the largest expression variability across all tumors (i.e., highly variable genes or HVG) were identified using the scran R package (version 1.12.1) (Lun et al. 2016), with FDR < 0.05. Manipulated data were analyzed using HCPC.

### Cell clustering using randomly-selected genes

Iterative HCPC analyses were performed on decreasing numbers of genes randomly selected among all detected genes. Analyses of ten distinct sets of randomly selected genes were performed for each size of gene sets (n = 5000, 4000, 2000, 1000, 500, 250, 100, 50, 25, 10 and 5). In these analyses, the number of clusters was set to the number of patient tumors in the dataset considered (number of clusters = number of tumors).

### Normalized Mutual Information (NMI) value calculation

The NMI value was used to evaluate the overall contribution of each tumor to each cluster of malignant cells. NMI was also used to evaluate the contribution of lineage subtypes to each cluster of non-cancerous cells hereafter designated as either normal when coming from non-pathological tissues or as non-malignant when coming from tumors. The NMI metric provides an external clustering validation that compares clustering results to a known truth (here tumor or lineage subtype labels) (Manning et al. 2008). A NMI value of 1 denotes that clusters gather objects (cells) corresponding to a single label, whereas a value of 0 denotes that all labels are split equally across all clusters (e.g. all tumors contribute equally to each cluster).

## Acknowledgments

This work was supported by grants from Région Ile-de-France (MSS fellowship), INCADGOS-Inserm_12560: SiRIC CURAMUS (financially supported by the French National Cancer Institute, the French Ministry of Solidarity and Health and Inserm), and La Fondation pour la Recherche Médicale - Equipes FRM 2020. We are grateful to Aurélien Barré (Bordeaux Bioinformatics Center, France) for his help with recovery of datasets. We are highly grateful to all authors who made freely available the scRNA-seq datasets they developed.

## Author contributions

Conceptualization, MSS, EEH, CA, MPJ, HC; Methodology, MSS, LB, YE, EEH, MPJ; Computational analyses, MSS, YE, LB; Writing – Original Draft, MSS, MPJ; – Review & Editing, all; Supervision, MPJ; Project Administration, HC; Funding Acquisition, HC, MPJ, CA, YE.

## Figure legends

**Supplemental Table S1.** Lists of single-cell RNA-seq datasets analyzed.

**Supplemental Table S2.** List of NMI values corresponding to each data manipulation presented in Fig. 2.

**Supplemental File 1**. R scripts.

**Supplemental File 2.** Python scripts.

**Supplementary Fig. S1.**
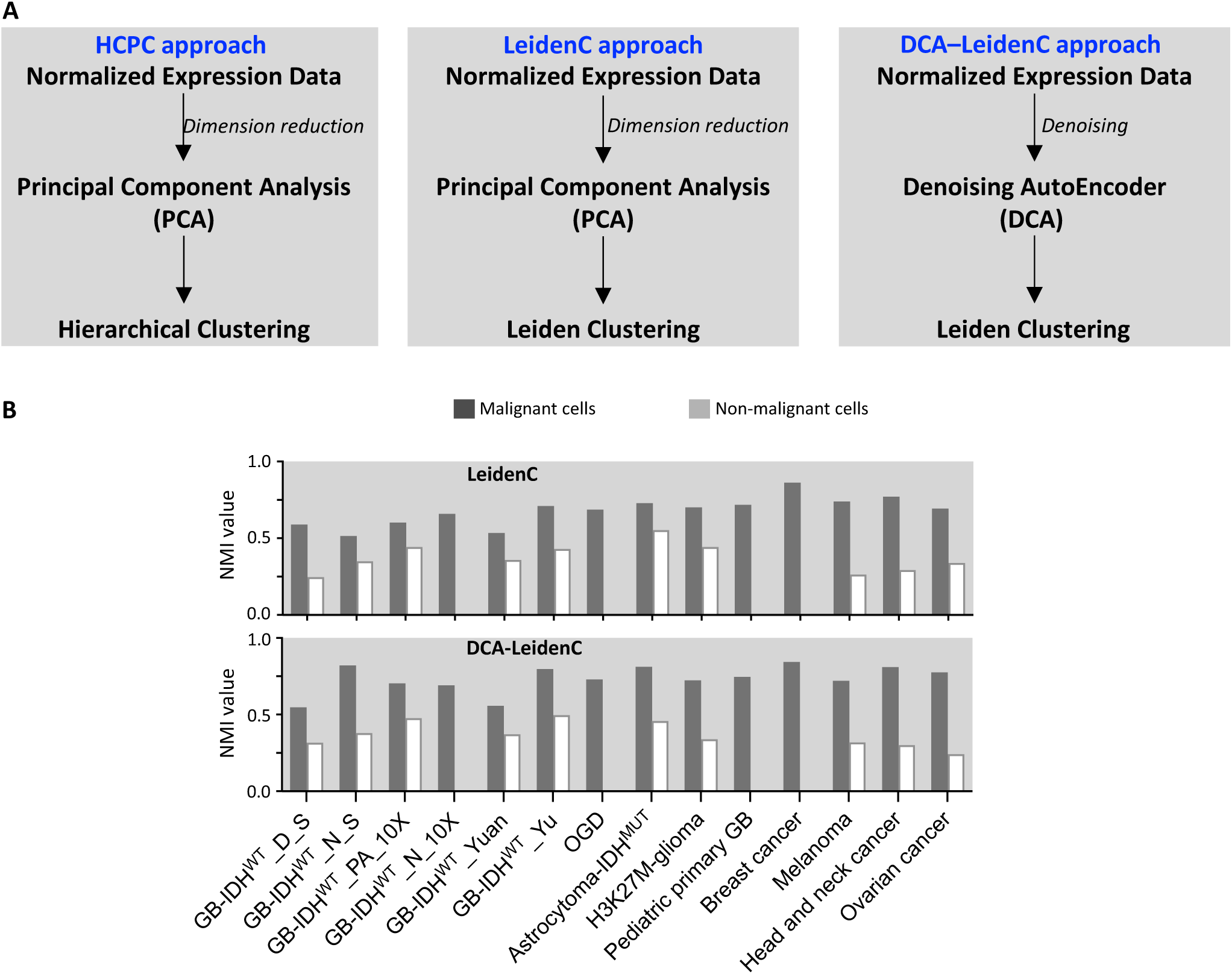
Analytical approaches used for cell clustering. Related to Figure 2. **A.** Overview of main differences between the three analytical approaches tested. **B.** Tumor contribution to cell clusters. Bar plot representations of Normalized Mutual Information (NMI) values corresponding to the grouping with LeidenC and DCA-LeidenC approaches based on expression of all detected genes of malignant, and when available non-malignant cells. GB: glioblastoma with wild-type form of the Isocitrate dehydrogenase (IDH) genes. IDH1^MUT^: mutant form of IDH1 gene. OGD: IDH1^MUT^ 1p/19q-codeleted Oligodendroglioma.

**Supplementary Fig. S2.**
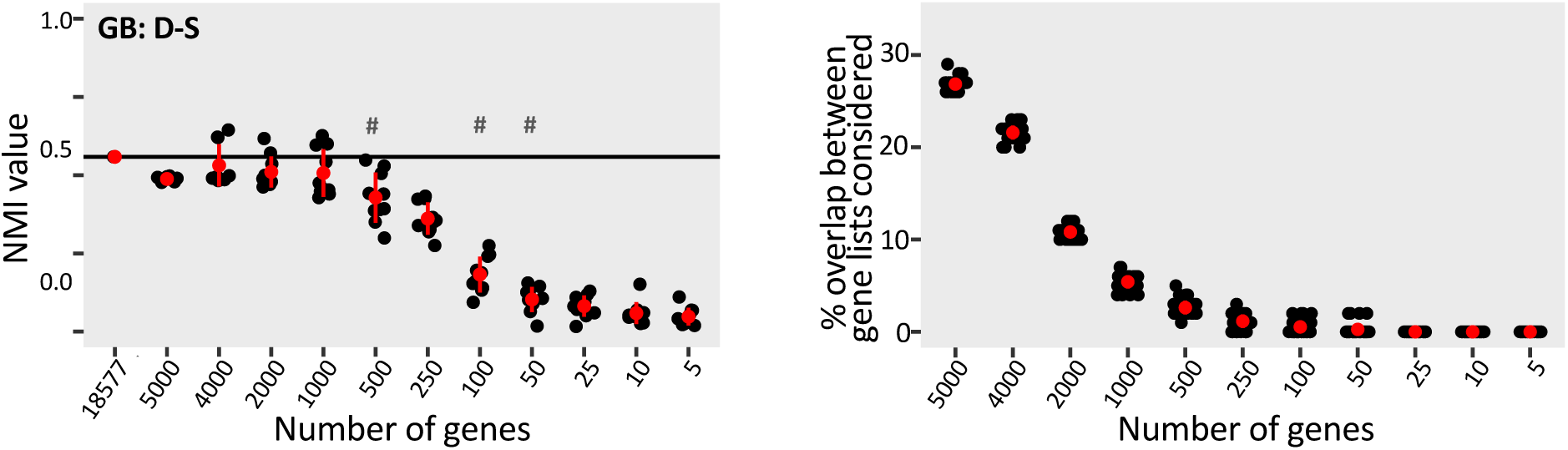
Combined expression of over 100 genes is sufficient to retrieve the identity of malignant cells of each glioblastoma tumor from the Darmanis-S dataset. Related to Figure 3. Left panel: Decreased NMI values with inclusion of decreased number of randomly selected genes in the grouping analyses. 10 independent analyses performed with randomly-selected genes for each gene number analyzed. Mean ± SD shown in red. #: statistically significant differences with the immediately prior sampling, Holm-Sidak’s multiple comparison test, adj-p value<0.05. Right Panel: overlap between gene lists used in analyses depicted in left panel.

**Supplementary Fig. S3.**
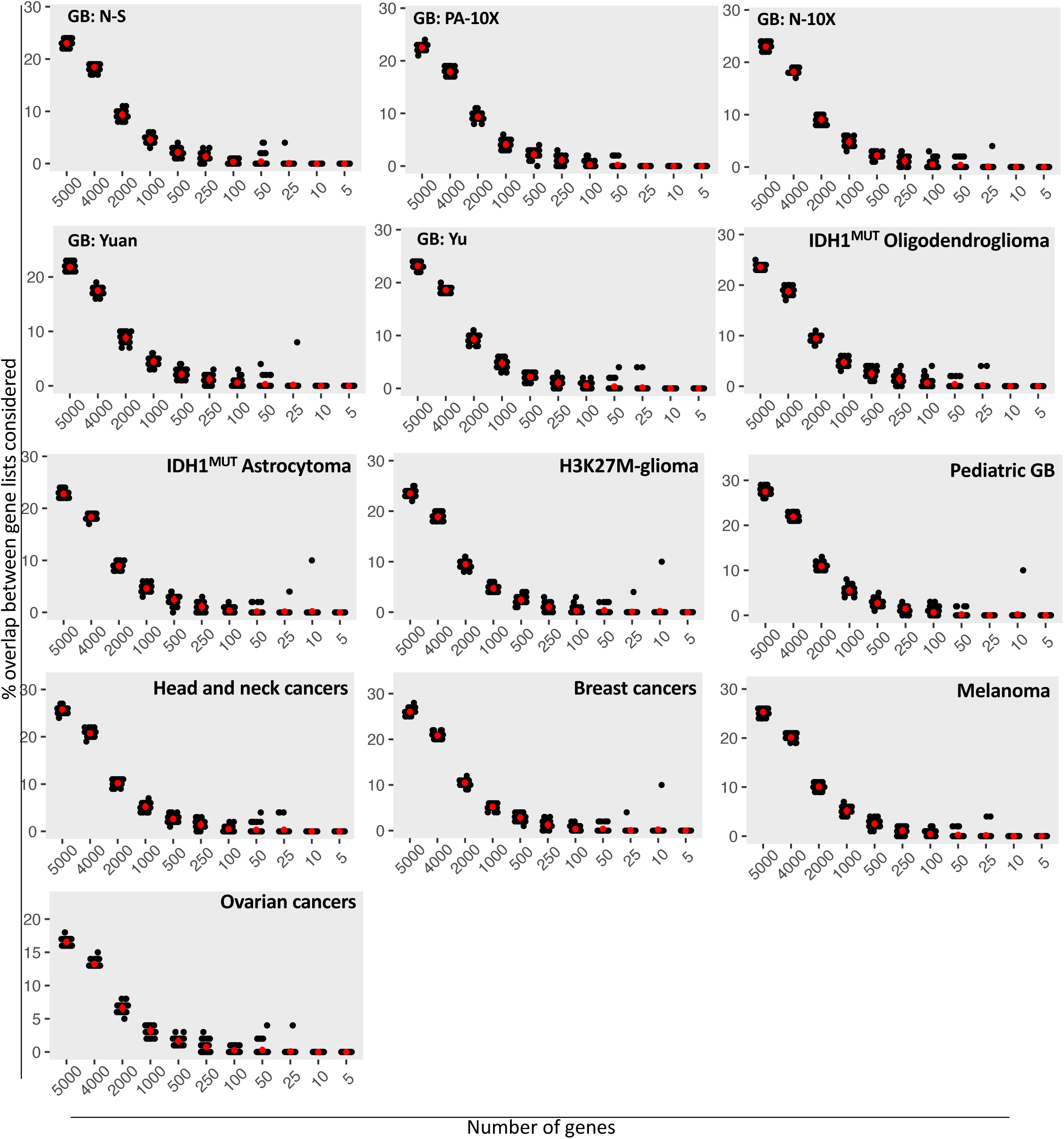
Overlap between lists of randomly-selected genes considered for grouping malignant cells from tumor samples studied. Related to Figure 3. Gene lists considered in down-sampling analyses of malignant cells are dissimilar from one another, with <30% overlap for gene sets containing 250-5000 genes and almost no overlap for gene set sizes ≤100. Mean ± SD in red.

**Supplementary Fig. S4:**
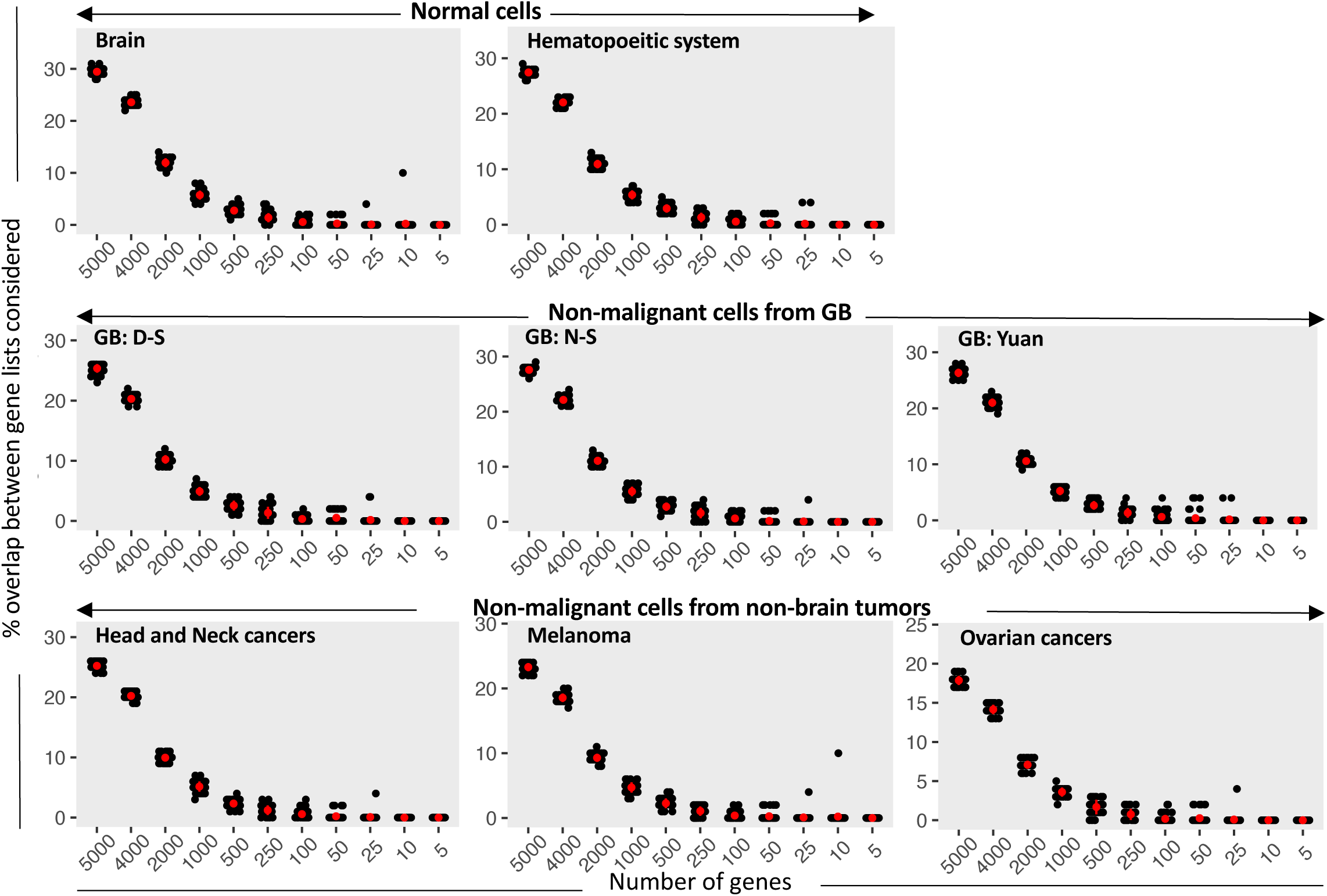
Overlap between lists of randomly-selected genes considered for grouping normal or non-malignant cells. Related to Figure 4. Gene lists considered in down-sampling analyses of non-malignant cells are dissimilar from one another, with <30% overlap for gene sets containing 250-5000 genes and almost no overlap for gene set sizes ≤100. Mean ± SD in red.

**Supplementary Fig. S5.**
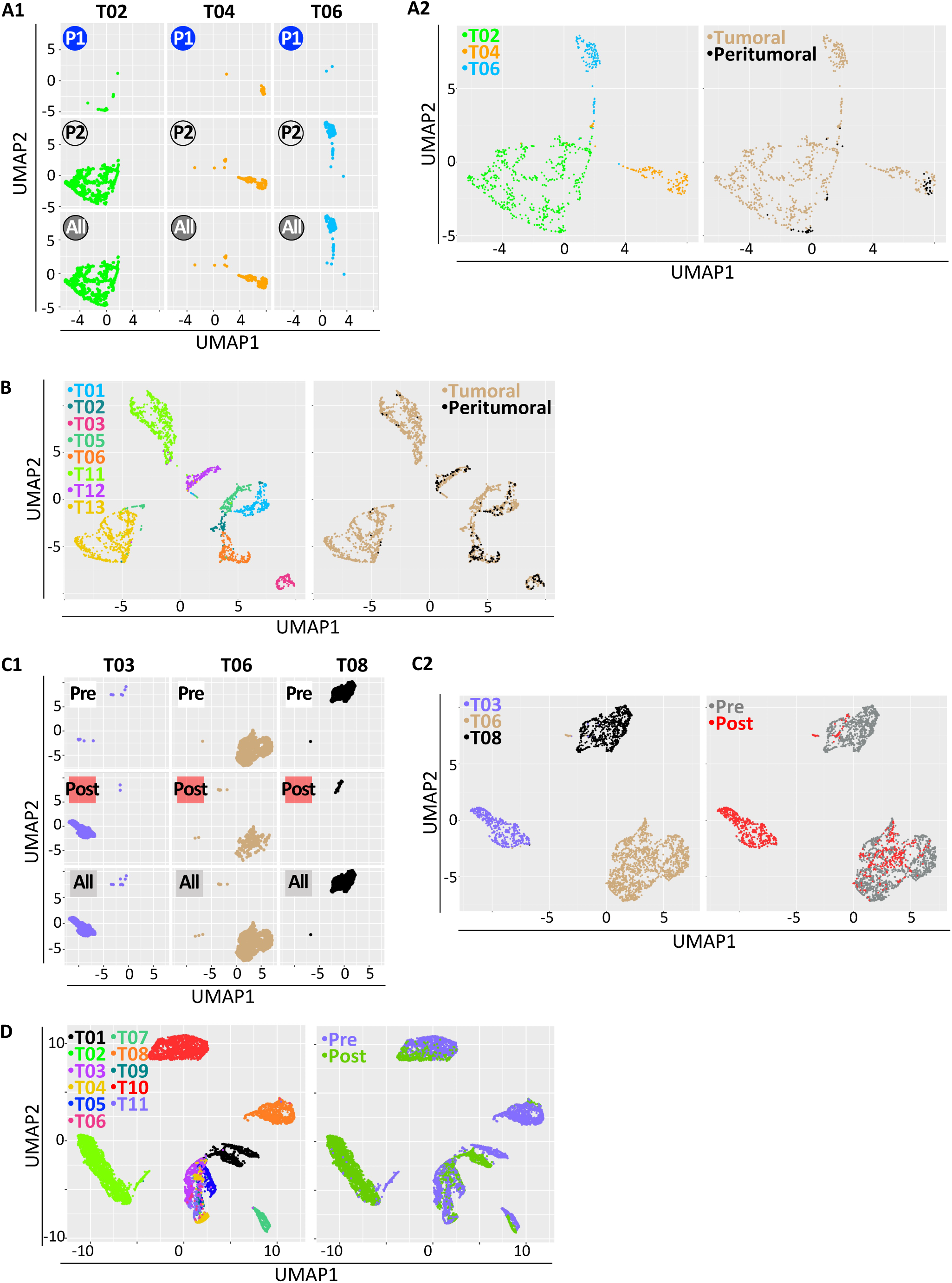
Tumor-specific identity of malignant cells is maintained through differing spatial environments and across time. Related to Figure 5. UMAP representations. **A.** Malignant cells collected from different glioblastoma regions. SMART-seq2 dataset from Darmanis et al 2017. **A1.** Results presented tumor by tumor. The numbers of sampled regions are circled. Blue circles correspond to malignant cells sampled from the peri-tumoral brain, clear circles to cells from the tumor, and grey circles to cells from all regions. **A2.** Results presented with all tumors. Cells colored by patient tumor (left panel) or tumor region (right panel). **B.** Malignant cell grouping of all sampled glioblastoma regions. GB scRNA-seq dataset from Yu et al 2020. Results detailed tumor by tumor in Fig. 5A presented here with all tumors on the graphs. Cells colored by tumor of origin (left panel) and tumor region (right panel). **C.** Malignant cells from basal cell carcinoma before or after treatment group according to the patient tumor from which they are harvested. 10X genomics dataset from Yost et al 2019. **C1.** Results presented tumor by tumor. White squares correspond to cells collected prior treatment, red squares to cells collected post-treatment and grey squares to cells from both time points. **C2**. Results presented with all tumors. Cells colored by patient tumor (left panel) or treatment time point (right panel). **D.** Malignant cell grouping of ovarian cancers collected at different treatment time points. Results detailed tumor by tumor in Fig. 5B presented here with all tumors on the graphs. Cells colored by tumor of origin (left panel) and treatment time point (right panel). scRNA-seq dataset from Zhang et al et al 2022.

**Supplementary Fig. S6.**
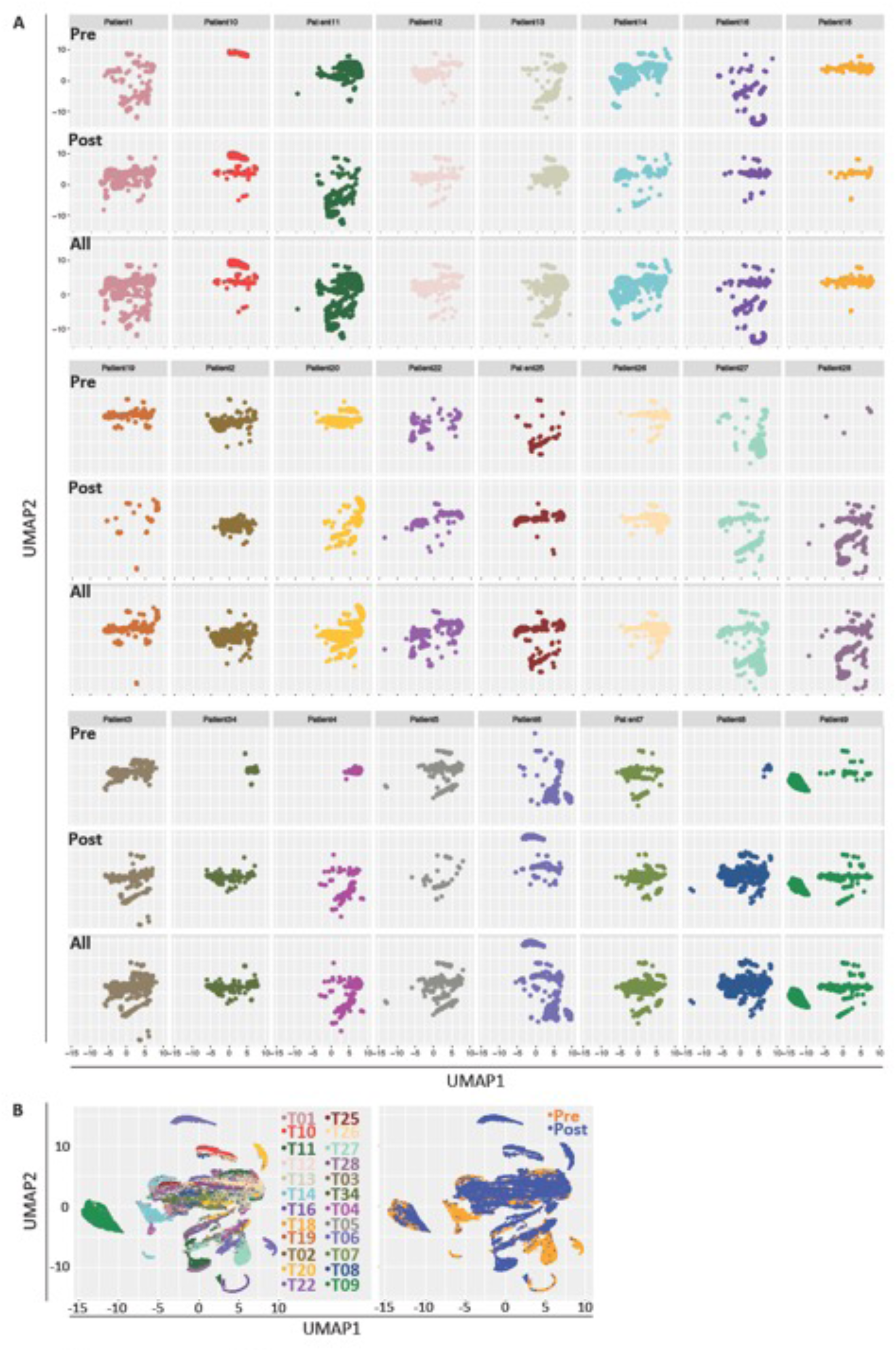
Tumor-specific identity of malignant cells is maintained over time. Related to Figure 5. UMAP representations. Malignant cells from the primary tumor of glioblastoma patients (Pre) or from the tumor recurrence after treatment (Post), group according to the patient tumor from which they are harvested. 10X genomics dataset from Wang et al 2022. **A.** Results presented tumor by tumor. **B**. Results presented with all tumors. Cells colored by patient tumor (left panel) or treatment time point (right panel).

**Supplementary Fig. S7.**
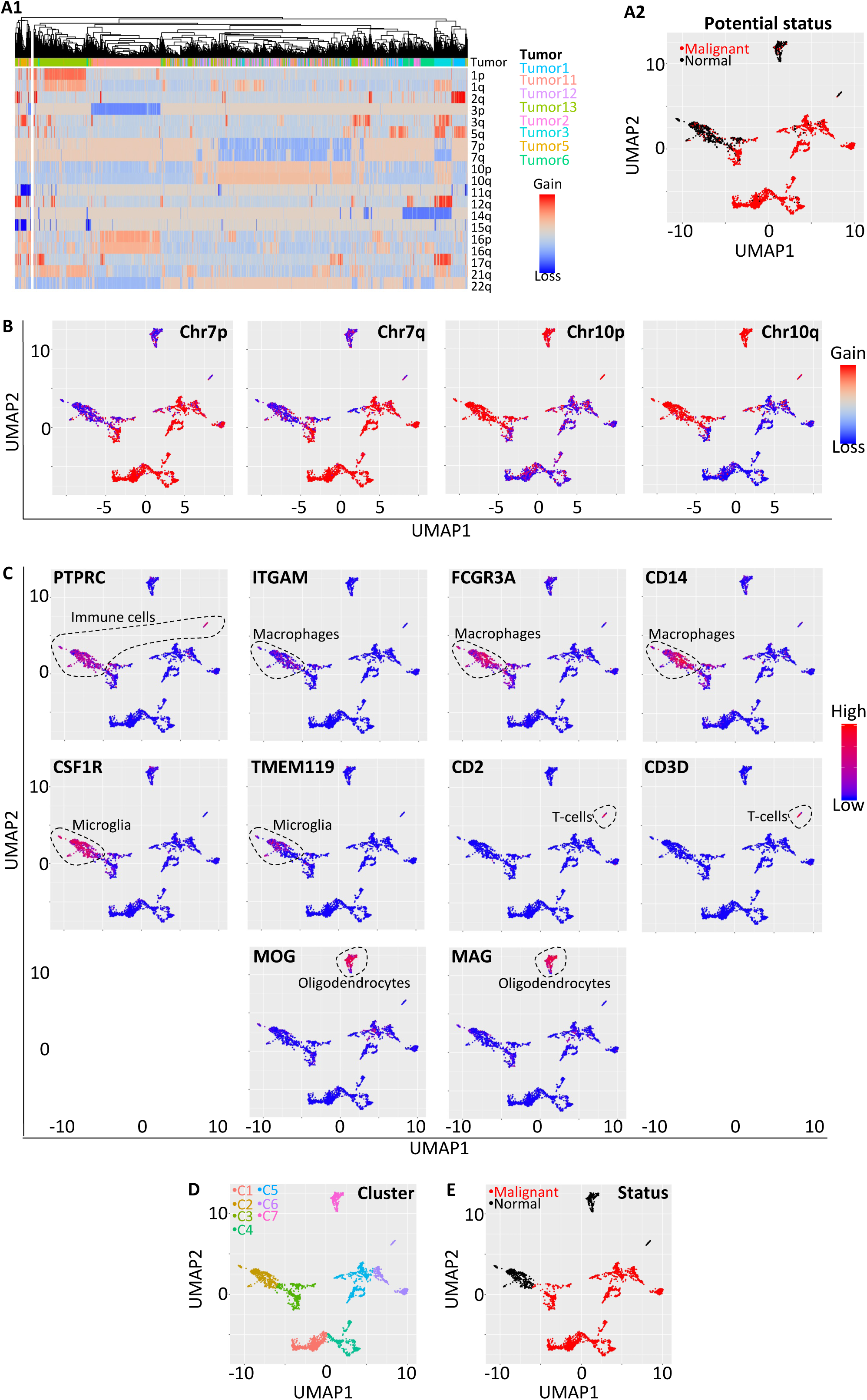
Identification of malignant and non-malignant cells in glioblastoma tumors from Yu et al 2020. Related to Methods.

